# Gene functional networks and autism spectrum characteristics in young people with intellectual disability

**DOI:** 10.1101/2020.05.11.088740

**Authors:** Diandra Brkić, Elise Ng-Cordell, Sinéad O’Brien, Gaia Scerif, Duncan Astle, Kate Baker

**Affiliations:** MRC Cognition and Brain Sciences Unit, University of Cambridge, 15 Chaucer Road, Cambridge, CB2 7EF, UK; Department of Experimental Psychology, University of Oxford, Anna Watts Building, Radcliffe Observatory Quarter, Woodstock Road, Oxford, OX2 6GG, UK

**Author notes:** contributed equally to this work. Corresponding author:* Kate Baker.

## Abstract

**Background:** Genes associated with Intellectual disability (ID) can be grouped into networks according to gene function. This study asked whether individuals with ID show differences in autism spectrum characteristics (ASC), depending on the functional network membership of their rare, pathogenic *de novo* genetic variants.

**Methods:** Children and young people with ID of known genetic origin were allocated to two broad functional network groups: synaptic physiology (n=29) or chromatin regulation (n=23). We applied principle components analysis to the Social Responsiveness Scale to map the structure of ASC in this population, and identified three components – Inflexibility, Social Understanding and Social Motivation. We then used Akaike Information Criterion (AIC) to test the best fitting models for predicting ASC components, including demographic factors (age, gender), non-ASC behavioural factors (global adaptive function, anxiety, hyperactivity, inattention) and gene functional networks.

**Results:** We found that, when other factors are accounted for, the chromatin regulation group showed higher levels of Inflexibility. We also observed contrasting predictors of ASC within each network group. Within the chromatin regulation group, Social Understanding was associated with inattention, and Social Motivation was predicted by hyperactivity. Within the synaptic group, Social Understanding was associated with hyperactivity, and Social Motivation was linked to anxiety.

**Conclusion:** We report that gene functional networks can predict Inflexibility, but not other ASC dimensions. Contrasting behavioural associations within each group suggests network-specific developmental pathways from genomic variation to autism. Simple classification of neurodevelopmental disorder genes as high risk or low risk for autism is unlikely to be valid or useful.

## Introduction

Intellectual disability (ID, defined as IQ <70 plus impaired adaptive function) and autism spectrum disorder (ASD, defined as persistent deficits in social communication and social interaction plus restricted, repetitive behaviours, interests, or activities) frequently co-occur, but are not synonymous (1). Understanding autism within the ID population is important, because autism predicts the complexity of educational, occupational and social support needs (2), and influences the well-being of family carers (3). One factor which can influence behavioural phenotypes, including autism, is the aetiology of each individual’s ID. At least 60% of individuals with severe ID have an underlying genetic diagnosis, which can now be readily diagnosed (4). However, the relationship between genetic diagnoses and autism is hotly contested. Some large cohort studies have presented evidence for “autism-predominant” neurodevelopmental disorder genes (5,6), whereas others argue strongly against this classification on both theoretical and empirical grounds (7). To resolve this question, systematic phenotyping is required to determine whether the genetic cause of ID predicts autism spectrum characteristics (ASC). However, addressing this question on a gene-by-gene basis holds several methodological challenges. Firstly, the rarity of each genetic disorder means that knowledge of phenotypic spectra can be skewed by small case numbers, not taking into account the expected variation in phenotypes within small groups, and rarely comparing across aetiologies associated with similar levels of ID severity. Secondly, cohort studies have typically relied on primary ascertainment diagnosis, or retrospective coding from medical notes, rather than acquiring standardised data. Thirdly, existing studies mainly focus on the presence or absence of categorical ASD diagnosis, rather than recognising that the characteristics contributing to ASD are diverse and vary along continuous dimensions such as social communication and repetitive behaviours (8,9). Previous studies of well-known syndromes associated with ASD, for example Fragile X Syndrome and Tuberous Sclerosis Complex, have highlighted considerable variation in atypical social behaviours contributing to ASD, and different predictors of ASD within each syndrome group (10,11). In essence, to understand autism in the context of ID-associated genetic disorders it is necessary to move beyond categorical diagnosis to investigate diverse social behavioural profiles and their underlying correlates.

In the current study, we apply novel strategies to investigate the relationships between genetic aetiology and dimensional ASC in young people with ID. Our first strategy is to reduce ascertainment bias by recruiting individuals after genetic diagnosis, irrespective of primary indication for genetic testing. Our second strategy is to collect standardised carer-report phenotyping assessments, appropriate for individuals with ID. Thirdly, we take a data-driven approach to analyses, by mapping the component structure of ASCs at single item level, then modelling predictors within the sample. Fourthly, we adopt a functional network phenotyping approach, meaning that we group participants according to known molecular and cellular functions of genetic variants, to detect convergent influences on behavioural outcomes, and provide insights into cognitive and neural mechanisms linking genetic cause to behavioural outcome (12). The current study compares two functional networks - a narrowly defined group of chromatin structural modifiers (components and regulators of the SWI/SNF chromatin remodelling complex), and a broader group encompassing direct and indirect modifiers of synaptic physiology. Chromatin modelling is essential for the establishment and maintenance of gene expression profiles to support neuronal differentiation, structural brain organisation, and flexibility of neuronal circuitry for learning (13,14). Synaptic transmission, its upstream regulation and downstream signalling, are fundamental to dynamic neurophysiological processes supporting perception, memory and action (15). Our hypothesis was that these functional networks could be associated with different dimensional autism characteristics, reflecting distinct underlying mechanisms.

## Methods

### Recruitment

The authors assert that all procedures contributing to this work comply with the ethical standards of the relevant national and institutional committees on human experimentation and with the Helsinki Declaration of 1975, as revised in 2008. All procedures involving human participants were approved by the NHS / HRA NRES Committee, East of England (11/EE/0330). Participants had been clinically identified as having neurodevelopmental impairments (developmental delay, intellectual disability, or behavioural difficulties) and referred for diagnostic genetic testing via clinical or research pathways. A pathogenic or likely pathogenic variant had been evaluated by local clinical geneticist as a causal or contributory factor for the individuals’ neurodevelopmental presentation. Information about the current study was provided to eligible participants’ families via regional genetics services, other clinical services, other research projects, family support groups, and via the project website. Parents of children under 16 gave written informed consent on behalf of their child. For participants with ID over the age of 16 lacking capacity to consent, a consultee was appointed.

### Group definitions

Functional networks groups (FNGs) were manually curated based on biochemical function, synaptic proteomics, GeneOntology (biological class), and PubMed searching (Supplementary Material Table 1). FNGs comprised 1) genes involved in chromatin structural regulation (“Chromatin” group), and 2) genes involved in synaptic transmission, synapse-associated cytoskeleton or post-synaptic intracellular signalling (“Synaptic” group). Participants in the study had variants in 15 different genes: 23 participants had variants in one of five Chromatin genes *(ARID1B, SETD5, EHMT1, KAT6B, SMARCA2),* and 29 participants with variants in one of ten Synaptic genes *(CASK, CTNNB1, DDX3X, DLG3, DYRK1A, PAK3, SHANK3, STXBP1, TRIO, ZDHHC9)* (Supplementary Material Table 2).

### Questionnaire and Interview Measures

Parents or carers completed the Vineland Adaptive Behaviour Scales, Second Edition, Survey Interview Form (Vineland; 16), Social Responsiveness Scale, Second edition (SRS; 17), Developmental Behaviour Checklist (DBC; 18), and Conners Parent Rating Scales (CPRS; 19).

### Data Analysis

We first addressed whether autism in this study population is best conceptualised as a unidimensional or multidimensional construct, via principal components analysis (PCA) of SRS items. In line with previous studies (8), component solution was selected on: 1) scree plots / percentage of variance explained, and 2) conceptual interpretability. We applied orthogonal rotation (Varimax with Kaiser normalization) to identify potentially diverging dimensions and underlying mechanisms. SRS total and dimension scores were normally distributed, and simple group comparisons were conducted via independent samples t-tests. To identify predictors of ASC dimensions we applied Akaike’s Information Criterion (AIC) modelling, corrected for small sample sizes (AICc). Information criteria modelling approaches allow inference from more than one model, controlling for over-dispersion and taking into account goodness of fit (20), when the true model is too complex to be estimated parametrically (21,22). AIC models included all participants with complete questionnaire and interview data (N = 45). A consistent set of potential predictors were included across all analyses: age, gender, global ability (Vineland Adaptive Behavior Composite), inattention (CPRS inattention subscale), hyperactivity (CPRS hyperactivity subscale) and anxiety (DBC anxiety subscale). Analyses comprised two steps: 1) a model selection step, geared to identify the best fitting models based on AICc values, with the most parsimonious models (i.e., lowest AIC value) favoured; 2) a multi-modal inference step, geared to infer the weight of individual predictors relative to the others, and the associated confidence intervals. Interaction terms were included within the models, to assess whether predictors were the same or different between groups. Analyses were performed using *glmulti* package in R (23).

## Results

### Sample characteristics

Demographics and descriptive data are displayed in Table 1. Fifty-two individuals (30 female) took part in the study. Groups were well-matched in gender and age. The Chromatin group had higher levels of global adaptive ability than the Synaptic group. The parents of 8 participants reported that their child had received a clinical diagnosis of ASD, pervasive developmental disorder not otherwise specified, or atypical autism, evenly distributed across FNGs. Groups did not differ in total SRS score, or % above cut-off for possible clinical diagnosis of ASD (X^2^ = .547, p = .46). Groups also did not differ significantly in non-ASC emotional and behavioural scores (DBC total, DBC anxiety subscale, Conners-3 inattention and hyperactivity subscales).

**Table 1.** Demographic and behavioural characteristics

|  | Chromatin (N = 23) |  | Synaptic (N = 29) |  | Independent samples t-test |
| --- | --- | --- | --- | --- | --- |
|  | Mean (SD) | Range | Mean (SD) | Range | *Chromatin-Synaptic* |
| Gender | 11F:12M | - | 19F:10M | - | - |
| Age | 12.93 (5.14) | 5 to 25 | 15.38 (5.23) | 7 to 26 | $t(50) = -1.69, p = .097, d = .47$ |
| Vineland Composite | 64.96 (11.90) | 41 to 96 | 49.59 (13.97) | 20 to 79 | $t(50) = 4.20, p < .001, d = 1.18$ |
| SRS Total (T) | 76.30 (12.73) | 53 to 96 | 75.52 (11.30) | 49 to 98 | $t(50) = .24, p = .814, d = .06$ |
| SRS raw score above 60; cut-off for clinical significance | 19/23 (82.60%) | - | 26/29 (89.66%) | - | - |
| CPRS Inattention (T) <sup>a</sup> | 79.71 (11.49) | 60 to 90 | 80.50 (10.99) | 55 to 90 | $t(43) = -.23, p = .816, d = .07$ |
| CPRS Hyperactivity (T) <sup>a</sup> | 69.05 (14.55) | 40 to 90 | 76.92 (15.03) | 42 to 90 | $t(43) = -1.78, p = .082, d = .53$ |
| DBC Total (percentile) | 61.65 (28.59) | 2 to 100 | 67.93 (26.49) | 18 to 98 | $t(50) = -.82, p = .416, d = .23$ |
| DBC Anxiety (percentile) | 55.39 (28.09) | 10 to 98 | 56.55 (25.51) | 10 to 98 | $t(50) = -.16, p = .877, d = .04$ |
| Factor 1 (Inflexibility) | .23 (1.06) | -1.52 to 2.20 | -.18 (.93) | -1.89 to 1.58 | $t(50) = -1.49, p = .143, d = .41$ |
| Factor 2 (Social Understanding) | -.17 (1.13) | -2.39 to 1.90 | .13 (.88) | -2.13 to 1.45 | $t(50) = -1.09, p = .280, d = .30$ |
| Factor 3 (Social Motivation) | -.03 (1.11) | -1.67 to 2.03 | .02 (.92) | -1.42 to 1.71 | $t(50) = -.167, p = .868, d = .05$ |

<sup>a</sup> 45 of 52 participants (21 in the Chromatin Group and 24 in the Synaptic Group) completed the CPRS.

### Mapping the structure of autistic behaviours in ID

First, PCA was run on all 65 items of the SRS-2. On visual inspection of the scree plot, there was a steep drop-off in the variance explained between three (37.1% variance explained) and four (41.4% variance explained) components (Supplementary Material 4). Solutions containing one, two, three, four, and five components were examined conceptually. Again, a three-component solution appeared to be optimal: with four- and five-component solutions, similar items were split into overlapping components, whereas with one- and two-component solutions, many items within the components were not aligned. Conceptually, our solution bears similarity to the model proposed by Nelson et al. (9), who explored the factor structure of SRS teacher-reported scores in children with autism and cognitive impairments. Based on these combined findings, a three-component solution was selected. In a second step, items with communalities < 0.4 (35 items) were excluded to maximise the overall communalities. The remaining 30 items were subjected to a second PCA. The full rotated component matrix for the three-component solution is displayed in Supplementary Material 5. Items showing cross-loading were included. In the final model, the KMO value was 0.597, and Bartlett’s test was significant (p <.001), i.e. sampling adequacy and data structure were appropriate for PCA with this reduced number of items. The model accounted for 51.87% of variance in item scores. Component 1 (Inflexibility) accounted for 22.62% of the variance in SRS item scores and includes items related to behavioural and cognitive flexibility, as well as ritualistic or compulsive behaviour (e.g. difficulty with changes to routine, fixated patterns of thought, or sensory sensitivity). Component 2 (Social Understanding) accounted for 19.69% of variance, and pertains to social awareness and cognition (e.g. knowing when invading others’ personal space, offering comfort to others when they are sad, or understanding cause and effect relations between events). Component 3 (Social Motivation) accounted for 9.56% of variance, and includes items related to disinhibition or withdrawal in social situations (e.g. avoiding starting interactions with others, avoiding emotional closeness with others, or having poor self-confidence in social settings). Mean component scores did not differ significantly between groups (Table 1). As a secondary analysis, we applied oblique rotation (Promax), and findings converged with primary analyses (Supplementary Material 6).

### Whole sample predictors of ASC components

For each ASC component, three top-ranked AIC models with goodness of fit indices (AIC weights, deviance, and ΔAIC) are provided in Table 2. For Inflexibility, the top-ranked model had an AIC weight of 0.323, or 32% of probability of being the best model. For Social Understanding, the top-ranked model AIC weight was 0.202. For Social Motivation, the top-ranked model AIC weight was 0.339. There were multiple models competing for the top rank (ΔAIC<2; see Table 2 and Supplementary Material 7). To reduce model uncertainty, model averaging and parameter estimation were calculated for each ASC dimension (Figure 1). This indicated that the most important predictors of Inflexibility were anxiety, hyperactivity, and genetic group (FNG). Higher Inflexibility was associated with higher levels of hyperactivity and anxiety, and being in the Chromatin group. For Social Understanding, likely predictors of impairment were lower global adaptive ability and elevated hyperactivity. For Social Motivation, only hyperactivity was predictive across the sample, with lower levels of hyperactivity associated with social withdrawal.

**Figure 1.**
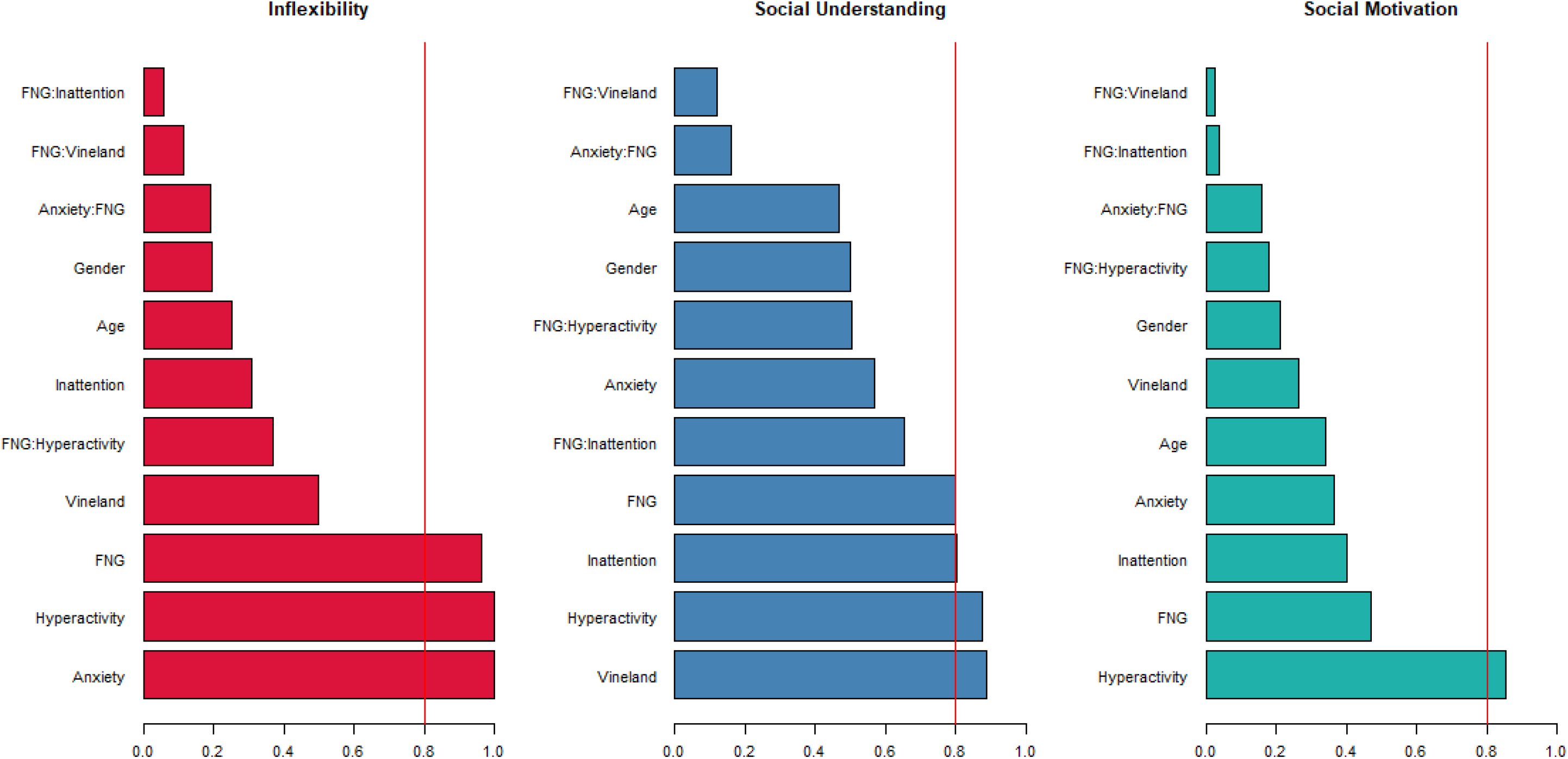
Relative importance of predictors for each ASC component, averaged across the set of candidate models. The arbitrary threshold of .08 was applied as cut-off for the most relevant predictors. FNG=Functional Network Group (Synaptic or Chromatin)

**Table 2.**
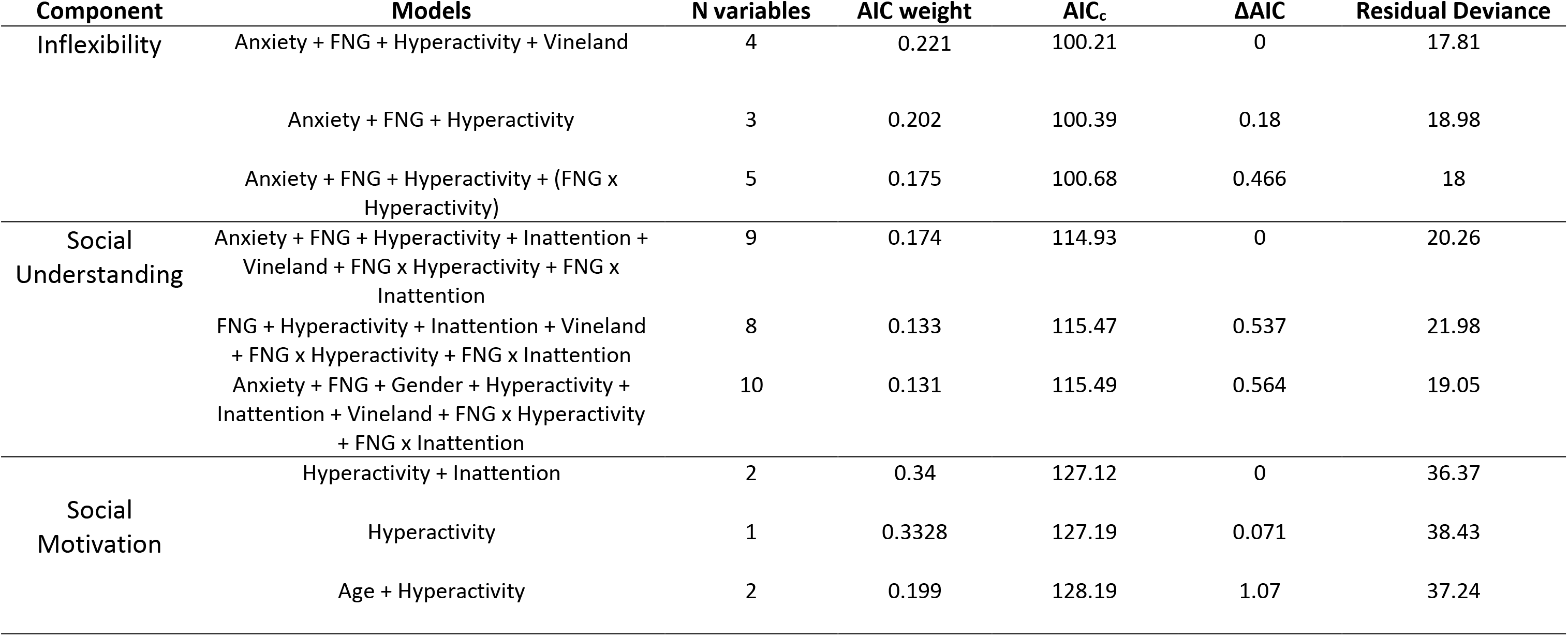
AIC Models for Autism Spectrum Characteristics, within Whole Sample

### Within-group predictors of ASC components

Figure 2 illustrates associations between ASC components and behavioural predictors within both functional network groups. The positive effect of global adaptive function on Social Understanding was the same across both groups, and no group-specific effect of global adaptive function was observed for either Inflexibility or Social Motivation. We observed an interaction between group, hyperactivity and Inflexibility, whereby the association between hyperactivity and Inflexibility is more pronounced within the Chromatin group (whereas the association between anxiety and Inflexibility is constant across groups). We also observed group-specific predictors of impaired Social Understanding (anxiety and inattention for the Chromatin group, hyperactivity for the Synaptic group). For Social Motivation, contrasting relationships were observed within groups: hyperactivity predicts social disinhibition within the Chromatin group, whereas anxiety predicts social withdrawal within the Synaptic group. For effect sizes of each coefficient and predictor, see Supplementary Material 8.

**Figure 2.**
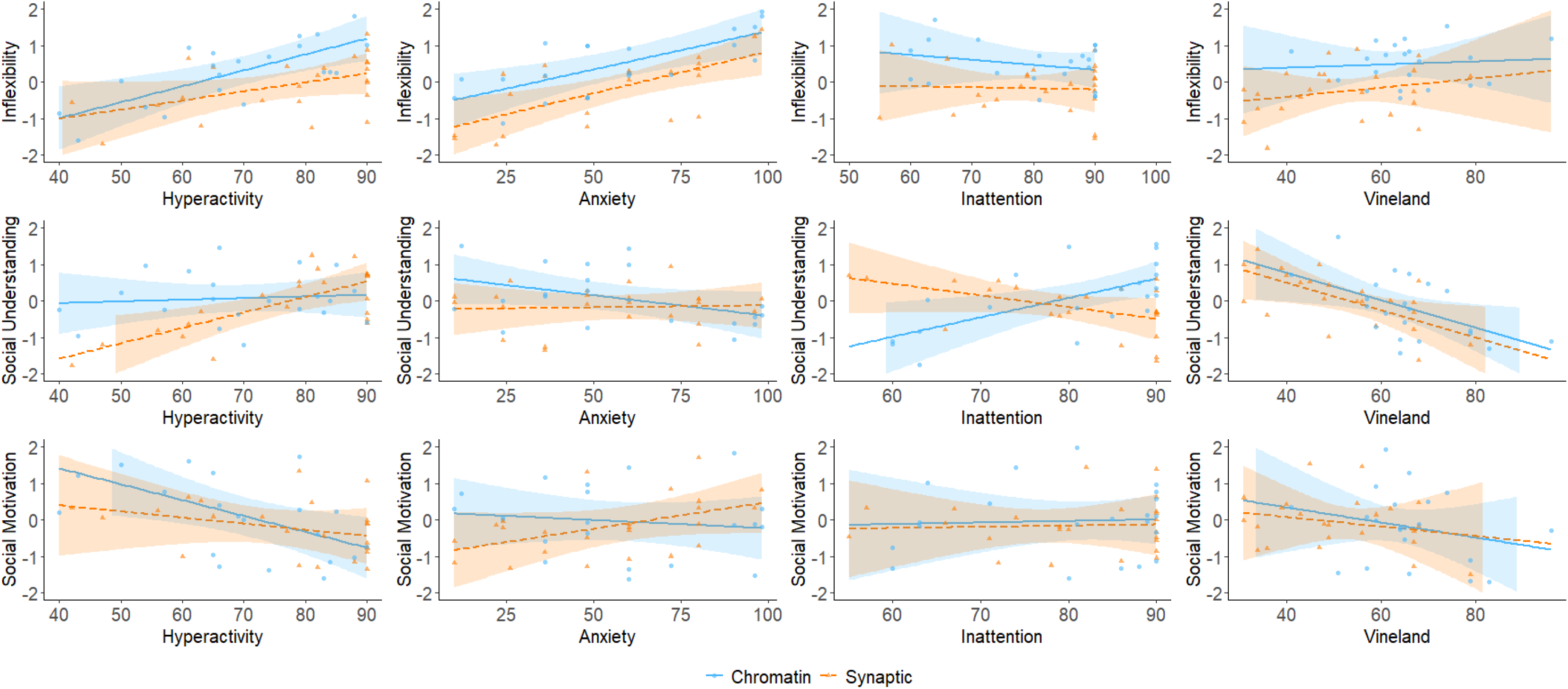
Relationships between ASC and non-ASC behavioural characteristics, within Functional Network Groups.

## Discussion

Numerous data-driven analyses have implicated discrete functional networks such as chromatin regulation, synaptic communication and cytoskeletal architecture in the neuronal origins of ASD (24). However, to date there has been no evidence that pathogenic variants within these gene sets influence the prevalence or types of autistic characteristics amongst individuals with ID. In this study, we address this hypothesis directly. The overall likelihood of autism characteristics was high across the sample and did not differ between groups. However, after separating autism-relevant questionnaire items into dimensions, and taking background variables into account, we found that gene functional networks predicted specific aspects of autism phenotype, and predicted co-occurrence between ASC and other behavioural characteristics.

### Disorders of chromatin regulation, inflexibility and cognitive control

Our within-sample modelling found that disorders of chromatin regulation are associated with elevated Inflexibility. These specific behavioural characteristics can have important knock-on consequences for individuals’ access to educational and psychosocial interventions, and exert strong influence on family life and well-being. Inflexibility may be masked by more overt difficulties, for example communication impairments or oppositional behaviour. Our results indicate that professionals should have heightened awareness of Inflexibility and its consequences when assessing and supporting individuals with Chromatin-related disorders. An important next step is to consider how chromatin regulation is related to Inflexibility, at the levels of cognitive development, neural systems and molecular neurobiology. The observed relationship between hyperactivity and inflexibility within the Chromatin group suggests disproportionate impact of chromatin dysregulation on cognitive control systems. We also found that hyperactivity and inattention predicted social disinhibition and social understanding, uniquely within the Chromatin group, further supporting the potential importance of cognitive control for social development of these individuals. At a neural level, cognitive control relies upon functional integration between multiple cortical areas. Chromatin-associated genes could influence functional integration via early development of relevant cortical and subcortical structures, later white matter development, or dynamic remodelling of neural networks (25)26).

### Disorders of synaptic physiology and social-emotional development

Individuals within the Synaptic group had more severe ID on average, which did not translate to higher SRS total scores or simple differences in ASC factor scores, emphasising that autism characteristics are not an inevitable consequence of global cognitive impairments. The predictors of ASC dimensions within the Synaptic group contrast with those observed within the Chromatin group, suggesting that a distinct set of developmental mechanisms may contribute to the social-emotional difficulties of the Synaptic group. Within this group (only), we observed that anxiety and social withdrawal are correlated, and hyperactivity and social understanding are negatively linked. Further investigation is warranted to determine whether these associations highlight specific relationships between synaptic physiology, motor control, emotional arousal and social interaction, or are common associations amongst individuals with severe ID. For the Social Motivation dimension, higher rates of residual deviance and lower model weights indicate that there are unmeasured predictors contributing to variability in this heterogeneous component. Further research should disentangle the factors contributing to social withdrawal versus social disinhibition, both of which can be distressing and impairing for the individual and their social circle.

### Limitations

The functional networks approach is advantageous in identifying broad group-based associations and spotlighting potential mechanistic convergence, however we openly acknowledge that the approach will mask potentially important gene-specific characteristics. Our approach of allocating genes to network groups is based on integration of multiple literature sources, each limited by existing functional data. Boundaries between networks are difficult to define; for example, chromatin-associated genes will have downstream effects on synaptogenesis, neurotransmission and plasticity by regulating expression of synaptic-relevant targets (27–29). We included components of the Wnt signalling pathway *(DDX3X* and *CTNNB1)* in the “Synaptic” group, because of emerging evidence that Wnt signalling directly “tunes” neurotransmitter release and modulates synaptic plasticity (30). Similarly, we included *DYRK1a* in the Synaptic group because there are multiple lines of experimental evidence supporting a direct role for this kinase in regulation of presynaptic vesicle cycling (31). Ultimately both data-driven and experimental approaches to functional network definitions would avoid bias in group allocations. Several further limitations are recognised. First, our sample size was small. The study was intended to be exploratory, and future pre-registered, multi-site studies with larger samples should test the stability of our three-component solution, explore a wider range of potential predictors (e.g. epilepsy, motor deficits, sensory impairments), and determine the robustness of the functional networks phenotyping approach and our specific findings. Larger samples would allow parallel PCA to determine whether ASC structure is constant across FNGs. This study deployed carer-report questionnaire measures only, and future studies could obtain richer insights via multi-informant reports, interview schedules, observational methods and neuropsychological assessments. ASC are expected to change with chronological and developmental age, perhaps in a gender-modified fashion (32), necessitating longitudinal studies. Lastly, socioeconomic status and family characteristics such as household structure, parental education, family stress and parental mental health may also interact with ASC, with complex bidirectional relationships between child and family factors (33), which may also encompass genetic diagnosis (3).

### Conclusions

In this study, the genetic cause of an individual’s ID (classified by functional network) did not predict overall likelihood of autistic features, but did influence dimensional autism characteristics and co-occurrences. These results indicate that dimensional phenotyping and data-driven modelling can enhance the prognostic utility and clinical relevance of genetic diagnosis for individuals with ID. Chromatin regulator variants were associated with elevated Inflexibility, suggesting disproportionate impact on neural systems underlying cognitive control. Furthermore, we report early insights into multiple pathways contributing to Social Understanding and Social Motivation, which may be differentially influenced by gene functional network groups. These data highlight the diversity of social and emotional characteristics that contribute to autism in the context of ID, and corresponding diversity of genetic and neurodevelopmental mechanisms. Future research should seek to replicate and extend these findings, and investigate the molecular, neural, cognitive and interpersonal mechanisms contributing to the emergent tapestry of social function for individuals with ID and their families.

## Acknowledgments

This study would not have been possible without the time and dedication of study participants, their families, and carers. We thank the clinicians, diagnostic laboratory scientists and research personnel at UK Regional Genetic Centres, the DDD study and the IMAGINE-ID study, who identified potential participants and publicised this study. The DDD study presents independent research commissioned by the Health Innovation Challenge Fund [grant number HICF-1009-003] a parallel funding partnership between the Wellcome Trust and the Department of Health, and the Wellcome Trust Sanger Institute [grant no. WT098051]. See Nature 2015; 519:223-8 or www.ddduk.org/access.html for full acknowledgement. The IMAGINE-ID study (http://imagine-id.org/) is funded by UK Medical Research Council and Medical Research Foundation (IMAGINE-ID, grant number MR-N022572-1). The research team acknowledges the support of the National Institute for Health Research, through the Comprehensive Clinical Research Network and Rare Genetic Disease Research Consortium.

## Author contribution

ENC and SOB acquired the data. DB and ENC analysed and interpreted the data. DB, ENC and KB drafted the paper and all other authors revised it. DA, GS and KB conceived and designed the study. All authors read and approved the final manuscript.

## Data availability

The data that support the findings of this study are available to other ethically-approved research projects from the corresponding author, KB.

