## Supplementary Material for "Gene functional networks and autism spectrum characteristics in young people with intellectual disability"

1. Evidence for functional network classification of chromatin-associated and synaptic-associated Intellectual Disability genes

2. Participant numbers by gene and functional network group

3. Within-sample predictors of ASC dimensions: model selection and multimodal inference

4. Extraction and rotation values for Principal Components Analysis

5. Rotated component matrix for three-component solution

6. PCA solution with oblique rotation

7. Complete table of top-ranked models with ΔAIC <2, for each ASC component

8. Effect size plots

### Supplementary Material Table 1. Evidence for functional network classification of chromatin-associated and synaptic-associated Intellectual Disability genes

| Gene | Protein | Biochemical and cellular function | Human adult brain expression | Human developmental brain expression | Synaptic proteome | Synaptic-relevant GO Biological processes | Chromatin-relevant GO Biological processes |
|---|---|---|---|---|---|---|---|
| Resource | ncbi.nlm.nih.gov/omim | genecards.org | braineac.org | hbatlas.org | SynaptomeDB.org | http://amigo.geneontology.org/ | http://amigo.geneontology.org/ |
| CHROMATIN GROUP | | | | | | | |
| ***ARID1B*** | At-Rich Interaction Domain-Containing Protein 1b | Component of SWI/SNF chromatin remodeling complexes, changing chromatin structure by altering DNA-histone contacts within a nucleosome in an ATP-dependent manner | Max – cerebellum<br>Min - thalamus | Peaks day 150, stable postnatal | NO | • Differentiation of interneurons<br>• excitatory / inhibitory balance<br>• Dendritic morphology | • SWI/SNF complex<br>• chromatin-mediated maintenance of transcription |
| ***EHMT1*** | Euchromatic Histone Methyltransferase 1 | Histone methyltransferase that specifically mono- and dimethylates 'Lys-9' of histone H3 (H3K9me1 and H3K9me2, respectively) in euchromatin | Max – white matter<br>Ubiq in cortex | Peaks early gestation, declines through prenatal, stable postnatal | NO | Nil | • DNA methylation<br>• chromatin organization |
| ***KAT6B*** | Lysine Acetyltransferase 6b | Histone acetyltransferase which may be involved in both positive and negative regulation of transcription. Required for RUNX2-dependent transcriptional activation | Not in database | Peaks day 100, stable postnatal | NO | Nil | • negative regulation of transcription, DNA-templated<br>• positive regulation of transcription by RNA polymerase II<br>• nucleosome assembly |
| ***SMARCA2*** | Swi/Snf-Related, Matrix-Associated, Actin-Dependent Regulator Of Chromatin, Subfamily A, Member 2 | Component of SWI/SNF chromatin remodeling complexes that carry out key enzymatic activities, changing chromatin structure by altering DNA-histone contacts | Max – cortex<br>Min – white matter | Increases during prenatal life. Stable postnatal | NO | Nil | |

| | | | | | | | |
|---|---|---|---|---|---|---|---|
| | | within a nucleosome in an ATP-dependent manner. | | | | | |
| ***SETD5*** | Set Domain-Containing Protein 5 | Displays histone methyltransferase activity and monomethylates 'Lys-9' of histone H3 in vitro. Probable transcriptional regulator that acts via the formation of large multiprotein complexes that modify and/or remodel the chromatin. | Max – cerebellum<br>Min - medulla | Peaks early gestation, steady decline | NO | Nil | • covalent chromatin modification<br>• regulation of chromatin organization |
| SYNAPTIC GROUP | | | | | | | |
| ***CTNNB1*** | Catenin, Beta-1 | Downstream component of the canonical Wnt signalling pathway | Max – cerebellum, thalamus<br>Min – putamen, SNIG | Peaks early gestation, then steady | YES | • synaptic vesicle transport<br>• synaptic vesicle clustering<br>• synaptic transmission<br>• Wnt signalling pathway, calcium modulating pathway | • Positive regulation of transcription |
| ***DDX3X*** | Dead/H Box 3, X-Linked | Multifunctional ATP-dependent RNA helicase. | Ubiq | Peaks early gestation, then steady | YES | • Wnt signalling pathway | • DNA helicase activity<br>• Translational and transcriptional regulation<br>• Positive regulation of gene expression<br>• RNA secondary structure unwinding |
| ***DLG3*** | Discs, Large Homolog 3 | Membrane-associated guanylate kinase | Max – hippocampus, cortex<br>Min – medulla, white matter | Increases across prenatal, declines from late childhood | YES | • structural constituent of postsynaptic density<br>• regulation of postsynaptic membrane neurotransmitter receptor levels<br>• maintenance of postsynaptic density structure<br>• regulation of NMDA receptor activity | nil |
| ***PAK3*** | p21 protein (Cdc42/Rac)-Activated Kinase 3 | Serine/threonine protein kinase. Acts as downstream effector of small GTPases | Max – hippocampus, cortex | Increases during prenatal life. Stable postnatal | NO (but PAK1 yes) | • synapse organization<br>• dendritic spine morphogenesis | Nil |

| | | | | | | | |
|---|---|---|---|---|---|---|---|
| | | | Min – cerebellum, white matter | | | | |
| ***SHANK3*** | Sh3 And Multiple Ankyrin Repeat Domains 3 | Major scaffold postsynaptic density protein | Max – hippocampus, putamen<br>Min – white matter, medulla | Not in database (SHANKS 1 and 2 – increases across prenatal, stable / declines postnatal) | YES | • synapse assembly<br>• positive regulation of long-term neuronal synaptic plasticity<br>• positive regulation of synaptic transmission, glutamatergic<br>• AMPA and NMDA glutamate receptor clustering | Nil |
| ***STXBP1*** | Syntaxin-Binding Protein 1 | Regulation of synaptic vesicle docking and fusion through interaction with GTP-binding proteins | Max – cortex<br>Min – white matter | Increases during prenatal life. Stable postnatal | YES | • vesicle docking involved in exocytosis<br>• regulation of synaptic vesicle priming<br>• negative regulation of synaptic transmission, GABAergic<br>• positive regulation of calcium ion-dependent exocytosis<br>• long-term synaptic depression | nil |
| ***TRIO*** | Triple Functional Domain | Guanine nucleotide exchange factor (GEF) for RHOA and RAC1 GTPases | Max – cerebellum, cortex<br>Min – white matter | Peaks day 150, declines postnatal | YES | • regulation of Rho protein signal transduction | Nil |
| ***ZDHHC9*** | Zinc Finger Dhhc Domain-Containing Protein 9 | Palmitoyltransferase | Max – white matter<br>Min - cerebellum | Peaks day 100, stable postnatal (adolescent increase?) | NO | • Protein targeting to membrane | Nil |
| ***DYRK1A*** | Dual-Specificity Tyrosine Phosphorylation-Regulated Kinase 1a | Dual-specificity kinase which possesses both serine/threonine and tyrosine kinase activities. Modulates alternative splicing by phosphorylating the splice factor SRSF6 | Max – cerebellum<br>Min – white matter | Peaks early gestation, declines through prenatal, stable postnatal | NO | • Protein tyrosine kinase | • negative regulation of mRNA splicing, via spliceosome<br>• transcription coactivator activity<br>• positive regulation of transcription, DNA-templated |

### Supplementary Material Table 2. Participant numbers by gene and functional network group

| Chromatin | | Synaptic | |
|---|---|---|---|
| **Gene** | **N** | **Gene** | **N** |
| *ARID1B* | 6 | *CASK* | 1 |
| *EHMT1* | 7 | *CTNNB1* | 1 |
| *KAT6B* | 1 | *DDX3X* | 9 |
| *SETD5* | 8 | *DLG3* | 2 |
| *SMARCA2* | 1 | *DYRK1A* | 2 |
| | | *PAK3* | 1 |
| | | *SHANK3* | 3 |
| | | *STXBP1* | 8 |
| | | *TRIO* | 1 |
| | | *ZDHHC9* | 1 |

### Supplementary Material 3. Within-sample predictors of ASC dimensions: model selection and multimodal inference

Step 1. *Model selection.* For each ASC component resulting from the PCA, we included the same set of variables: age, gender, global adaptive ability, FNG, and non-ASD behavioural traits (inattention, hyperactivity, and anxiety). Additionally, we explored interactions between genetic diagnosis (FNG) and predictors, to investigate shared vs distinctive associations. To do this we included in the model selection paradigm interaction terms between FNG and i) inattention, ii) hyperactivity, and iii) anxiety. AICc values were compared, with the most parsimonious models (i.e. lowest AIC value) favoured. The selection criteria for best fitting models, were based on *ΔAIC, or difference in AIC from between a model i and the first-ranked model (Johnson 2004.) Generally, models with ΔAIC<2 provide a substantially good fit to the data (Burnham and Anderson, 2002).*

Step 2. *Multimodal Inference*. In order to provide stable inference and parameter estimation, for each of the behavioural characteristics, we averaged across the top ranked models (*ΔAIC<2),* and computed the single coefficients' importance. The relative importance of the predictors or coefficients measures the relative likelihood that each predictor is part of the best model (Symonds 2011). This is estimated by summing the Akaike weights (ωAIC) across all the models in the candidate set. In short, the larger the weight is, the more important the variable or predictor, relative to the others. This procedure allows us to look at effects of closely related models, by measuring confidence intervals, and thus reducing model uncertainty (Johnson 2004).

### Supplementary Material 4. Scree plots, Extraction and rotation values for Principal Components Analysis

<u>Preliminary PCA on all SRS-2 items</u>

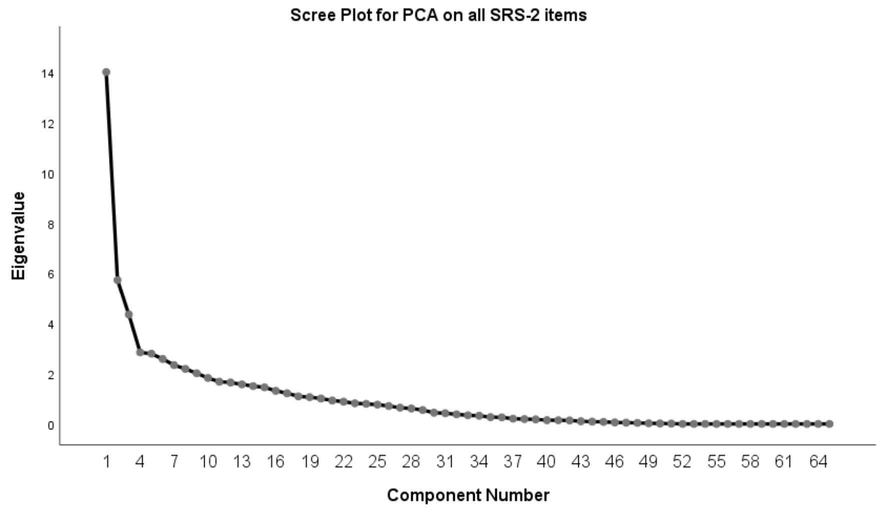


| | | Component | | | | |
|---|---|---|---|---|---|---|
| | | 1 | 2 | 3 | 4 | 5 |
| Extraction sums of squared loadings | Eigenvalue | 14.008 | 5.726 | 4.358 | 2.848 | 2.796 |
| | % of variance | 21.551 | 8.810 | 6.704 | 4.382 | 4.302 |
| | Cumulative % | 21.551 | 30.361 | 37.065 | 41.447 | 45.749 |

Extracted values for the first five components produced by initial PCA, performed on all 65 SRS-2 items.

**FNG ASC Dimensions – Supplementary Material**

Final PCA on reduced number of SRS-2 items (30 items)

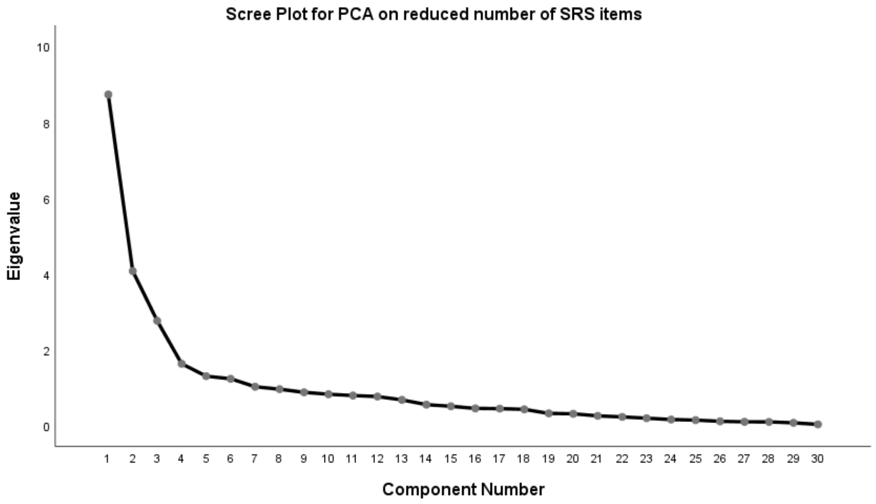


| | | Component | | |
|---|---|---|---|---|
| | | 1 | 2 | 3 |
| Extraction sums of squared loadings | Eigenvalue | 8.729 | 4.072 | 2.760 |
| | % of variance | 29.095 | 13.574 | 9.200 |
| | Cumulative % | 29.095 | 42.669 | 51.869 |
| Rotation sums of squared loadings | Eigenvalue | 6.786 | 5.908 | 2.867 |
| | % of variance | 22.618 | 19.694 | 9.556 |
| | Cumulative % | 22.618 | 42.312 | 51.869 |

Extracted and rotated values for the three component solution produced by a second PCA, performed on a reduced number of SRS-2 items (30)

### Supplementary Material 5. Rotated component matrix for three-component solution

| Item | Component | | |
|---|---|---|---|
| | **Inflexibility** | **Social Understanding** | **Social Motivation** |
| Difficulty with changes to routine | **.777** | .053 | .060 |
| Overwhelmed in situations with lots going on | **.776** | -.077 | .149 |
| Sensory sensitivity | **.752** | -.112 | -.058 |
| Has fixated patterns of thought | **.732** | .092 | .258 |
| Tense in social situations | **.731** | -.032 | .179 |
| Inflexible | **.697** | .172 | .158 |
| When stressed shows rigid behaviours | **.689** | .150 | .017 |
| Too literal | **.653** | .318 | .127 |
| Stares into space | **.634** | .183 | -.073 |
| Behaves in ways that are strange or bizarre | **.555** | **.516** | -.198 |
| Has difficulty relating to peers | **.530** | .371 | -.065 |
| Repetitive behaviours | **.451** | **.406** | **-.419** |
| Aware when being too loud | -.093 | **.706** | .046 |
| Aware of others' thoughts and feelings | .125 | **.695** | -.016 |
| Knows when standing too close to others | -.102 | **.693** | .003 |
| Offers comfort to others when they are sad | -.048 | **.686** | .258 |
| Understands cause and effect | .189 | **.675** | -.204 |
| Recognises when something is unfair | -.076 | **.658** | .156 |
| Understands the meaning of others' tone | .118 | **.613** | .227 |
| Regarded by others as odd | .292 | **.598** | -.060 |
| Awkward in turn-taking interactions | .289 | **.572** | .042 |
| Socially awkward | **.443** | **.550** | -.096 |
| Shows unusual sensory interests | **.447** | **.548** | -.178 |
| Difficulty communicating thoughts | .209 | **.542** | **.431** |
| Walks between people | .101 | **.505** | -.301 |
| Avoids initiating social interactions | .220 | -.091 | **.753** |

| Item | Component | | |
|---|---|---|---|
| | Inflexibility | Social Understanding | Social Motivation |
| Poo self-confidence | -.007 | .219 | **.700** |
| Avoids emotional closeness with others | .399 | .128 | **.601** |
| Silly | **.462** | .129 | **-.523** |
| Gets frustrated trying to communicate ideas | **.479** | -.008 | **.518** |

Variance explained: 51.87%. Extraction method: Principal Components with Varimax rotation and Kaiser normalization. Item loadings > 0.4 are in bold font.

### Supplementary Material 6. PCA solution with oblique rotation

*Pattern matrix for three-component solution with oblique rotation*

| Item | Component | | |
|---|---|---|---|
| | Inflexibility | Social Understanding | Social Motivation |
| Overwhelmed in situations with lots going on | **.839** | -.231 | .136 |
| Sensory sensitivity | **.820** | -.279 | -.073 |
| Difficulty with changes to routine | **.813** | -.100 | .049 |
| Tense in social situations | **.782** | -.172 | .168 |
| Has fixated patterns of thought | **.758** | -.035 | .250 |
| Inflexible | **.703** | .050 | .152 |
| When stressed shows rigid behaviours | **.698** | .018 | .152 |
| Stares into space | **.633** | .058 | -.079 |
| Too literal | **.627** | .212 | .125 |
| Has difficulty relating to peers | **.484** | .279 | -.064 |
| Behaves in ways that are strange or bizarre | **.479** | **.418** | -.195 |
| Aware when being too loud | -.245 | **.772** | .065 |
| Offers comfort to others when they are sad | -.192 | **.757** | .276 |
| Knows when standing too close to others | -.252 | **.757** | .021 |
| Recognises when something is unfair | -.216 | **.726** | .174 |
| Aware of others' thoughts and feelings | -.013 | **.711** | -.001 |
| Understands cause and effect | .059 | **.662** | -.192 |
| Understands the meaning of others' tone | -.001 | **.643** | .240 |
| Regarded by others as odd | .185 | **.570** | -.050 |
| Difficulty communicating thoughts | .110 | **.565** | **.443** |
| Awkward in turn-taking interactions | .188 | **.550** | .051 |
| Walks between people | .000 | **.493** | -.301 |
| Socially awkward | .354 | **.485** | -.090 |
| Shows unusual sensory interests | .359 | **.476** | -.172 |

| Item | Component | | |
|---|---|---|---|
| | **Inflexibility** | **Social Understanding** | **Social Motivation** |
| Avoids initiating social interactions | .256 | -.086 | **.750** |
| Poor self-confidence | -.050 | .286 | **.709** |
| Avoids emotional closeness with others | .399 | .097 | **.600** |
| Silly | **.460** | .003 | **-.529** |
| Gets frustrated trying to communicate ideas | **.512** | -.069 | **.512** |
| Repetitive behaviours | .391 | .307 | **-.418** |

Extraction method: Principal Components with Promax rotation and Kaiser normalization. Item loadings > 0.4 are in bold font.

**FNG ASC Dimensions – Supplementary Material**

*Pearson correlation matrix of component scores (Orthogonal and Oblique rotations)*

| | Inflexibility (Orthogonal) | | Social Understanding (Orthogonal) | | Social Motivation (Orthogonal) | | Inflexibility (Oblique) | | Social Understanding (Oblique) | | Social Motivation (Oblique) | |
|---|---|---|---|---|---|---|---|---|---|---|---|---|
| **N=52** | *r* | *p* | *r* | *p* | *r* | *p* | *r* | *p* | *r* | *p* | *r* | *p* |
| **Inflexibility (Orthogonal)** | | | .000 | 1.000 | .000 | 1.000 | .982 | <.001 | .193 | .171 | -.020 | .890 |
| **Social Understanding (Orthogonal)** | .000 | 1.000 | | | .000 | 1.000 | .191 | .176 | .981 | <.001 | -.073 | .606 |
| **Social Motivation (Orthogonal)** | .000 | 1.000 | .000 | 1.000 | | | .011 | .941 | -.020 | .886 | .997 | <.001 |
| **Inflexibility (Oblique)** | .982 | <.001 | .191 | .176 | .011 | .941 | | | .376 | .006 | -.023 | .873 |
| **Social Understanding (Oblique)** | .193 | .171 | .981 | <.001 | -.020 | .886 | .376 | .006 | | | -.096 | .499 |
| **Social Motivation (Oblique)** | -.020 | .890 | -.073 | .606 | .997 | <.011 | -.023 | .873 | -.096 | .499 | | |

**Supplementary Material 7. Complete table of top-ranked models with ΔAIC <2, for each ASD dimension.**

| Component | Models | N Variables | AIC Weight | AICc | ΔAIC | Residual Deviance | Explained Deviance or D squared |
|---|---|---|---|---|---|---|---|
| Inflexibility | Anxiety + FNG + Hyperactivity + Vineland | 4 | 0.221 | 100.21 | 0 | 17.81 | 0.623 |
| | Anxiety + FNG + Hyperactivity | 3 | 0.202 | 100.39 | 0.18 | 18.89 | 0.598 |
| | Anxiety + FNG + Hyperactivity + FNG x Hyperactivity | 5 | 0.175 | 100.68 | 0.466 | 18 | 0.619 |
| | Anxiety + FNG + Hyperactivity + Vineland + FNG x Hyperactivity | 7 | 0.136 | 101.19 | 0.973 | 17.1 | 0.638 |
| | Anxiety+FNG+Hyperactivity+Vineland+FNG x Vineland | 7 | 0.09 | 102.01 | 1.791 | 17.41 | 0.631 |
| | Age + Anxiety + FNG + Hyperactivity | 4 | 0.099 | 102.01 | 1.799 | 18.54 | 0.607 |
| | Anxiety + FNG + Hyperactivity + Inattention | 5 | 0.096 | 102.06 | 1.847 | 18.56 | 0.607 |
| Social Understanding | Anxiety + FNG + Hyperactivity + Inattention + Vineland + FNG x Hyperactivity + FNG x Inattention | 9 | 0.174 | 114.93 | 0 | 20.26 | 0.5952544 |
| | FNG + Hyperactivity + Inattention + Vineland + FNG x Hyperactivity + FNG x Inattention | 8 | 0.133 | 115.47 | 0.537 | 21.98 | 0.5607631 |
| | Anxiety + FNG + Gender + Hyperactivity + Inattention + Vineland + FNG x Hyperactivity + FNG x Inattention | 10 | 0.131 | 115.49 | 0.564 | 19.05 | 0.6193666 |
| | Anxiety + FNG + Hyperactivity + Inattention + Vineland + Anxiety x FNG + FNG x Hyperactivity + FNG x Inattention | 10 | 0.106 | 115.93 | 0.998 | 19.24 | 0.6156723 |

**FNG ASC Dimensions – Supplementary Material**

| Component | Models | N Variables | AIC Weight | AICc | ΔAIC | Residual Deviance | Explained Deviance or D squared |
|---|---|---|---|---|---|---|---|
| Social Understanding | FNG + Gender + Hyperactivity + Inattention + Vineland + FNG x Hyperactivity + FNG x Inattention | 9 | 0.1 | 116.04 | 1.107 | 20.76 | 0.5851734 |
| | Age + FNG + Gender + Hyperactivity + Inattention + Vineland + FNG x Hyperactivity + FNG x Inattention | 10 | 0.1 | 116.04 | 1.112 | 19.28 | 0.614705 |
| | Anxiety + FNG + Gender + Hyperactivity + Inattention + Vineland + Anxiety x FNG + FNG x Hyperactivity + FNG x Inattention | 11 | 0.097 | 116.1 | 1.168 | 17.85 | 0.6433202 |
| | Age + Gender + Hyperactivity + Vineland | 6 | 0.088 | 116.29 | 1.361 | 25.46 | 0.4912235 |
| | Age + FNG + Hyperactivity + Inattention + Vineland + FNG x Hyperactivity + FNG x Inattention | 9 | 0.07 | 116.76 | 1.835 | 21.1 | 0.5784112 |
| Social Motivation | Hyperactivity + Inattention | 4 | 0.34 | 127.12 | 0 | 36.37 | 0.154765 |
| | Hyperactivity | 3 | 0.328 | 127.19 | 0.071 | 38.43 | 0.1067548 |
| | Age + Hyperactivity | 4 | 0.199 | 128.19 | 1.072 | 37.24 | 0.1343813 |
| | Age + Hyperactivity + Inattention | 5 | 0.133 | 128.99 | 1.871 | 35.83 | 0.1672165 |

N variables = number of parameters for each model, aic weight= is the probability of each model of being the best model, or relative evidence for each model. aicc=aic criterion of model selection, corrected for smaller sample size, δaic=aic difference between the best fitting model (equal to zero) and the second best one. residual deviance=distance between the data and the model. fng=functional network group refers to the synaptic/chromatin grouping.

### Supplementary Material 8. Effect sizes plots

The pictured lines are the confidence intervals of each coefficient, and the white dot is their beta value, FNG=Functional Network Group refers to the synaptic/chromatin grouping. **Left** This plot shows the averaged estimates of the most important predictors of inflexibility and the direction of the effects. Higher Inflexibility was positively correlated with higher hyperactivity, anxiety, and general ability (Vineland). **Middle** Averaged coefficients of best predictors explaining the associations with difficulties in social understanding. Higher difficulties in social understanding were negatively correlated with age and lower general ability (Vineland). **Right** Averaged coefficients, across the candidate set of models, and their direction of effect for Social Withdrawal dimension. Higher difficulties in Social Withdrawal were significantly increased by higher rates of anxiety in the synaptic group.

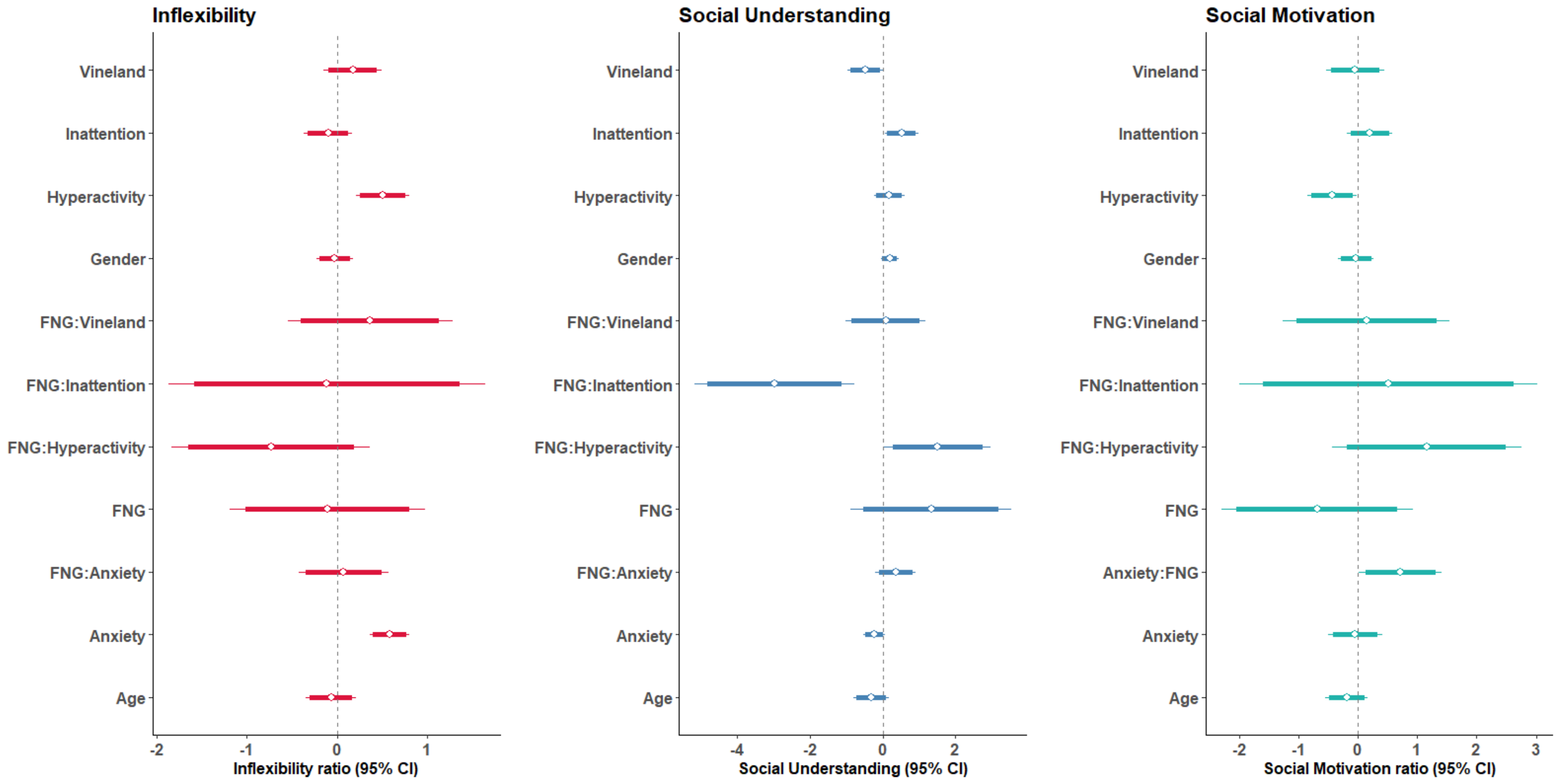
